# The computation of strategic learning in repeated social competitive interactions: Learning sophistication, reward attractor points and strategic asymmetry

**DOI:** 10.1101/346155

**Authors:** Thibaud Griessinger, Giorgio Coricelli, Mehdi Khamassi

**Affiliations:** TG: Laboratoire de neurosciences cognitives, Département d’études cognitives, École normale supérieure, INSERM, PSL Research University, 75005 Paris, France; GC: Department of Economics, University of Southern California, Los Angeles, USA; MK: Institute of Intelligent Systems and Robotics, Sorbonne Université, CNRS, Paris, France

## Abstract

Social interactions rely on our ability to learn and adjust our behavior to the behavior of others. Strategic games provide a useful framework to study the cognitive processes involved in the formation of beliefs about the others’ intentions and behavior, what we may call strategic theory of mind. Through the years, the growing field of behavioral economics provided evidence of a systematic departure of human’s behavior from the optimal game theoretical prescriptions. One hypothesis posits that human’s ability to accurately process the other’s behavior is somehow bounded. The question of what constraints the formation of sufficiently high order beliefs remained unanswered. We hypothesize that maximizing final earnings in a competitive repeated game setting, requires moving away from reward-based learning to engage in sophisticated belief-based learning. Overcoming the attraction of the immediate rewards by displaying a computationally costly type of learning might not be a strategy shared among all individuals. In this work, we manipulated the reward structure of the interaction so that the action displayed by the two types of learning becomes (respectively not) discriminable, giving a relative strategic (resp. dis) advantage to the participant given the role endorsed during the interaction. We employed a computational modeling approach to characterize the individual level of belief learning sophistication in three types of interactions (agent-agent, human-human and human-agent). The analysis of the participants’ choice behavior revealed that the strategic learning level drives the formation of more accurate beliefs and eventually leads to convergence towards game optimality (equilibrium). More specifically we show that the game structure interacts with the level of engagement in strategically sophisticated learning to explain the outcome of the interaction. This study provides the first evidence of a key implication of strategic learning heterogeneity in equilibrium departure and provides insight to explain the emergence of a leader-follower dynamics of choice.

**AUTHOR SUMMARY:** Dynamic interaction between individuals appears to be a cornerstone for understanding how humans grasp other minds. During a strategic interaction, in which the outcome of one’s action depends directly on what the other individual decides, it appears crucial to anticipate the other’s actions in order to adjust our own behavior. In theory, choosing optimally in a strategic setting requires that both players hold correct beliefs over their opponent’s behavior and best-respond to it. However, in practice humans systematically deviate from the game-theoretical (equilibrium), suggesting that our ability to form accurate beliefs is cognitively and/or contextually constrained. Previous studies using computational modelling suggested that during a repeated game interaction humans vary in the sophistication of their learning process leading to the formation of beliefs over their opponent’s behavior of different orders of complexity (level of recursive thinking such as “I think that you think that …”). In this work we show that the individual engagement in sophisticated (belief-based) learning drives the convergence towards equilibrium and ultimately performance. Moreover, we show that this effect is influenced by both the game environment and the cognitive capacity of the participants, shaping the very dynamic of the social interaction.

**DATA AVAILABILITY:** The authors confirm that upon publication the raw behavioral data and Matlab code for reconstruction of all figures, computational models and statistical analyses will be made available for download at the following URL: https://zenodo.org/

## I INTRODUCTION

Inferring someone’s intention is key to adapt to the behavior of others and maximize the outcome of social interactions [1]. It enables to establish shared action plans and efficient coordination in cases of cooperation [2]. It also enables anticipation of an opponent’s actions in cases of competition such as in strategic games.

Recently, an emphasis has been placed on one particular feature of this mind reading ability: using the past experience to predict the behavior of a conspecific [3]. Strategic interactions during repeated competitive games have been proven to be a useful experimental paradigm to capture the behavioral dynamics revealing such theory of mind in human and non-human primates [4], as they consist in social situations where one’s choice outcome critically depends upon the action of the other. Game-theory provides formal solutions to strategic interactions, modelled as games, through the concept of Nash Equilibrium (NE) and its refinements [5]. The so-called Mixed Strategy Nash Equilibrium (MSNE), for instance, prescribes a probability distribution over possible actions that ensures to each involved agent that they would have no incentive to deviate if they all follow it. In practice, however, humans typically deviate from equilibrium [6]. Moreover, the way human subjects progressively learn to reach MSNE is still little understood. One hypothesis, which encounters growing support, lies on the idea that convergence towards MSNE distribution during repeated games requires that both players, who aim to maximize their earnings, should learn and adjust beliefs through predictions over the opponent’s behavior in order to best-respond to it [13]. Nevertheless, the important variability observed in this process [15] makes the characterization of the underlying learning mechanism difficult.

Research in cognitive neuroscience suggests that, in probabilistic tasks, humans learn to adjust their decisions based on expected values computed from previously experienced outcomes (model-free reinforcement learning, RL) but also through the incorporation of (probabilistic) beliefs over the action-outcome contingencies underlying the structure of such environment (model-based RL) [19], [20]. Indeed in such tasks, reward convey at least two types of information, often correlated: the affective (hedonic) value embodied in the monetary reward and the information (predictive) value about the structure of the world [21]. The ability to use the latter to maintain and update, using prediction accuracy, a mental representation of the choice environment has been found to encompass social interactions as well [22], [23]. In fact, by pairing the normative framework of game theory to the computational approach in neurosciences, recent research has shown that during repeated games, humans can engage in model-based learning using an iterative computation of the strategic information provided by the interaction [24], [25]. Such strategically sophisticated computations drive higher order beliefs that incorporate the level of influence of one’s past actions on her opponent’s choice behavior, thus allowing for more accurate predictions [26]. Such hierarchy of belief computation has been observed at the brain level, with common brain structures implicated in model-based (non-social) and belief-based (social) learning computations (medial prefrontal cortex) [27], and higher-order belief (strategic) learning incorporating signals from areas involved in theory of mind (temporo parietal junction) [28].

Crucially, all these studies reported important heterogeneity in the level of engagement in belief-based learning, linked to variation in overall performance. However, none of the previous studies directly investigated the relationship between the human’s ability to engage in strategic learning and the observed deviation from best response distribution, and ultimately MSNE. Taken together these results yet suggest that humans’ propensity to follow optimality prescription from game-theory requires to disengage from reward-oriented, model-free, learning and fully engage in belief-based learning.

We hypothesized that, depending on how the reward structure interacts with the MSNE prescription in a repeated strategic game, human performance in the game may be differently affected so that it does not necessarily reflect an individual’s general level or ability of strategic learning. Previous studies suggest that the amplitude of the payoffs interferes with the propensity to follow the MSNE [7], and that the symmetric nature of a game might facilitate the belief formation over the opponent’s behavior through perspective taking [29], [30].

We developed a novel 2×2 competitive game setting, symmetric in payoff amplitude and expected payoff, so that the two players would earn the same if they both follow the MSNE distribution. The structure of the payoff matrix was however designed to lead to *strategic asymmetry* where one player’s highest rewarded action would happen to be, at the informational level, the one the MSNE prescribes to choose the most (advantageous role), while for the other player the attractive action (focal point) would be the one she should choose the less (disadvantageous position). If following the optimal distribution of choice is conditioned on the ability to consider the strategic structure beyond the payoffs value to engage in belief learning, then our game should lead to strategic asymmetry. We made the secondary hypothesis that in the repeated version of this game, humans with different individual strategic learning levels (SL) would differ in their capacity to overcome this asymmetry and lead to observable differences in the final earnings between the advantageous and disadvantageous roles in the game.

We ran 2 distinct experiments with the same game setting: In the first one human subjects play against each other, while in the second we specifically manipulate the level of subjects’ computerized opponent. Beforehand, we simulated agents interacting repeatedly through our competitive game, all modeled as simple learning algorithms varying from reward-based to sophisticated belief-based computations [25] and developed to capture different levels of strategic learning sophistication (SL). As anticipated we show that, at the population level, the game payoff matrix lead to a strong strategic asymmetry, such that the agents playing in the disadvantaged position see their loss reduced only when they engage in higher SL level than their opponent. We moreover found that the observed deviation from game optimality (MSNE) is mainly driven by the individual propensity to depart from reward-based learning and engage in sophisticated belief-learning, and that individuals in the disadvantageous position were driven by loss reduction and constrained by their own SL learning capacity, while subjects in the advantageous position were mainly adjusting their best response to the estimated behavior of their opponent. Importantly for our computational hypothesis, behavioral results from both experiments matched the predictions made in simulation. Strikingly, only subjects endorsing the disadvantageous role (hence pressed towards their own limit) showed a SL level which was stable across opponent, as if the reduction of the strategic asymmetry was cognitively bounded [31]. These findings thus provide a possible explanation for the discrepancy between previous studies in which no correlation between SL level in strategic games and cognitive abilities was observed [24].

## II MATERIALS AND METHODS

### 1 Experiment 1: Computer against computer

The game is a two-by-two (two players, two actions) (payoffs) asymmetric game, with a unique Mixed strategy equilibrium (**Fig. 1A).** The expected payoffs at the mixed strategy Nash equilibrium are the same for both players.

**Figure 1.**
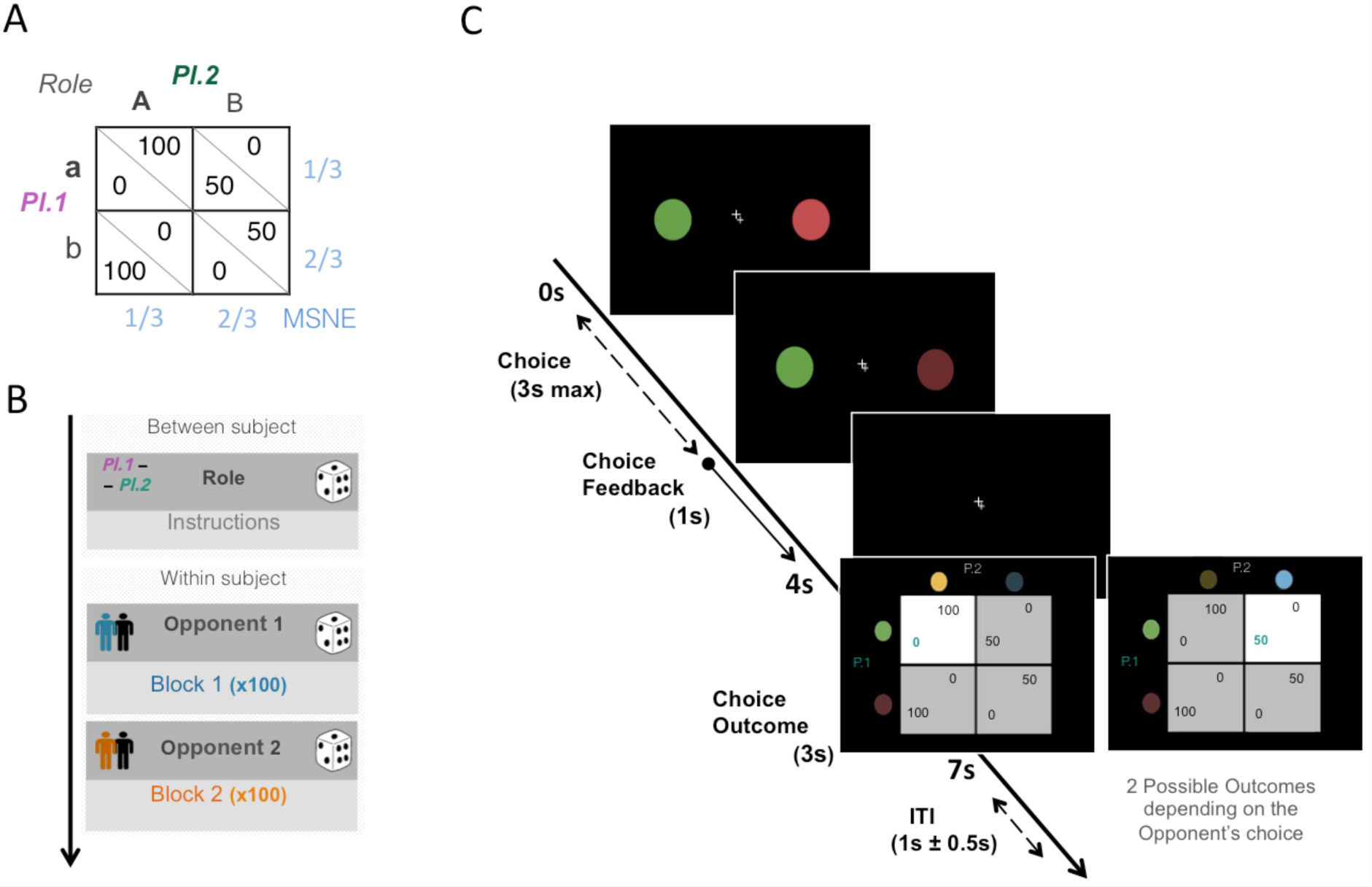
Experimental design of experiment 1. (A) Payoff matrix of the repeated game, in points. In light blue the action probabilities prescribed by the mixed strategy equilibrium (MSNE). (B) We manipulated 2 variables: the role (within subject level), and the opponent (between subject level). At start subjects were randomly assigned to one of the two roles in the game (player 1 or 2). After being instructed, they were randomly paired to an anonymous counterpart during a block of 100 repetitions of the game, and to a different counterpart during the second trial block. (C) Trial Structure: At each trial the two game actions, represented by the randomly assigned colors, were presented for 3s to each player. The choice was made by pressing the corresponding button (left or right). 4s after the trial onset, both players were simultaneously provided with the outcome feedback of their choice for 3s (the cell matrix matching to the 2 players choices was highlighted and the points won displayed in turquoise).

To make predictions about the effect of the strategic asymmetry of our payoff matrix on subjects’ behavior we simulated different computerized agents playing a repeated version of our game in each of the 2 roles, with different levels of strategic sophistication. Such a simulation analysis allows us to not only test the robustness of our design but also to make precise predictions regarding the effect the individual level of strategic learning would have on the dynamics of play of humans interacting through this experimental setting (see Text S1 for details on the computational modelling and the simulation analysis).

To mimic inter-individual variation of strategic learning we used 3 computational models varying in their level of strategic sophistication (SL): a simple reinforcement algorithm learning only from the outcomes obtained through its past choices; a fictitious play best responding to the probability of each opponent’s choice computed from its history of actions; and an Influence model, i.e. a 2nd order fictitious taking into account the influence of its own past choices in the computation of the opponent’s probability of play [25]. Each simulation consisted in 2 computerized agents, endorsing one of the 2 roles and modeled by one of the 3 models, playing against each other during 100 repetitions of the stage game.

### 2 Experiment 2: Human against human

#### a Population

64 participants (29 male, 35 female; ages 27.1±9.4) took part in the experiment. They were students at the University of Lyon 1, France, who had previously joined the recruitment system on a voluntary basis. These volunteers gave written informed consent for the project which was approved by the French National Ethical Committee. All participants were right-handed, medication-free, with normal eyesight, no history of neurological disorders.

#### b Experimental Design

The first experiment consisted in a repeated interaction against another anonymized participant. One of the 2 roles was randomly attributed to each participant at the beginning of the experiment. Each subject interacted with two different human opponents, one after the other in two trial blocks of 100 repetitions of the stage game with complete choice feedback (**Fig. 1B).** These two opponents were randomly selected among the participants assigned the opposite role to the subject. Points earned at each trial were accumulated through each block and summed up to determine their final payoff which would ultimately be converted to euros according to a predetermined rule. Each subject was initially instructed of the 2 stimuli representing her two available actions in the task, the payoff structure of the game, and trained to learn the stimulus-outcome contingencies of the payoff matrix. Each action was made of a different colored circle randomly picked from 4 possible colors (all controlled for luminance). The 4 colors were randomly assigned to each pair of subjects in the first block, and kept unchanged in the second interaction block (thus constraining the re-matching random procedure in the second block). At each trial, both subjects had 3 seconds to select one of the 2 colors displayed at the left and right of the screen (randomized order across trials), the chosen one was highlighted for 1 second as choice feedback. 4s after the trial onset, both players were simultaneously provided with the outcome feedback of their choice and the one of their opponent for 3s. The outcome feedback screen consisted in the payoff matrix (note that depending of the role endorsed in the game the matrix was flipped so that subjects were always presented as row player), with the cell corresponding to the matching of the 2 players choices highlighted and the points won by the subject displayed in turquoise (**Fig. 1C).** This display ensured minimal framing effect, while controlling for participants’ awareness of the underlying payoff structure of the game.

We also provided to the subjects an additional task which consisted of a series of four different types of 2×2 static (one-shot) games [16]. The goal was to test the endogeneous hypothesis of strategic learning sophistication developed in Griessinger & Coricelli [26]. We hypothesized that participants with a SL level in the repeated game (captured by our computational approach from the game behavior in the main task) would also display a higher strategic reasoning (SR, expressed as their capacity to conform to equilibrium play when a game is not repeated and no feedback is provided). All the subjects came back a second time to the lab a week later to complete a series of cognitive tasks. Both the additional experiment and cognitive tasks are detailed in Text S1.

#### c Computational modeling

To capture the level of strategic learning of the subjects we first used the computational approach introduced in the simulation analysis (Exp.1): 3 computational models, fcorresponding to 3 different levels of strategic sophistication, were fitted individually to each choice series from the two trial block independently (Q-Learning, Fictitious and Influence models). The underlying assumption is that the higher the level of strategic complexity of the model that best fits a subject’s behavior, the higher her strategic learning engagement in the interaction. As detailed in the **Text S1** we also tested additional models to control for the reliability of our computational approach.

### 3 Experiment 3: Human against computerized opponents

#### a Population

76 participants (36 male, 40 female; ages 18–30) took part in the experiment. They were student at the University of Trento, Italy, who had previously joined the Cognitive and Experimental Economics Laboratory (CEEL) recruitment system on a voluntary basis. All participants were right-handed, medication-free, with normal eyesight, no history of neurological disorders. The Ethics Commission of the University of Trento approved the experiment. Informed consent was obtained from each subject before the experiment. Data collection was performed blind to the conditions of the experiment.

#### b Experimental Design

The experimental design remained unchanged: participants were randomly assigned to one of the 2 roles of the same game, with the same trial structure and timing, and also played 2 blocks of 100 trials each. Nevertheless, this time they did not play against another randomly picked human participant, but rather played against 2 computerized learning agents: a fictitious play (low SL) and an Influence (High SL), one after the other in a random order. To be fully consistent with the previous experiment, we used as model parameters of the 2 opponents the average best fitting values obtained in the first experiment (details in Text S1).

All statistical analyses were performed using Matlab (www.mathworks.com) with the addition of the Statistical toolbox and other free-download functions. All stimuli and feedbacks were presented using PsychToolBox and appeared on a uniform black background.

## III RESULTS

### 1 Experiment 1

Our simulation results reveal an important advantage of Players 1 over Players 2 in our game. Not only simulated agents playing as Player 1 performed better than Players 2 in the game, but Players 2 had to be consistently higher SL level than their opponent in order to win more points **(Fig. 2).** To ensure that this game propriety does not depend on the tuning of our simulations we replicated the simulation analysis with different proxies of the SL level such as the parameter λ of the Influence model. This parameter captures the weight of the second order fictitious update in the computation of the opponent’s action probability. Our additional simulation analyses systematically show that the only way for Players 2 to outperform their opponent is to engage in a higher level of Strategic Learning **(Fig. S1 in Text S1).**

**Figure 2.**
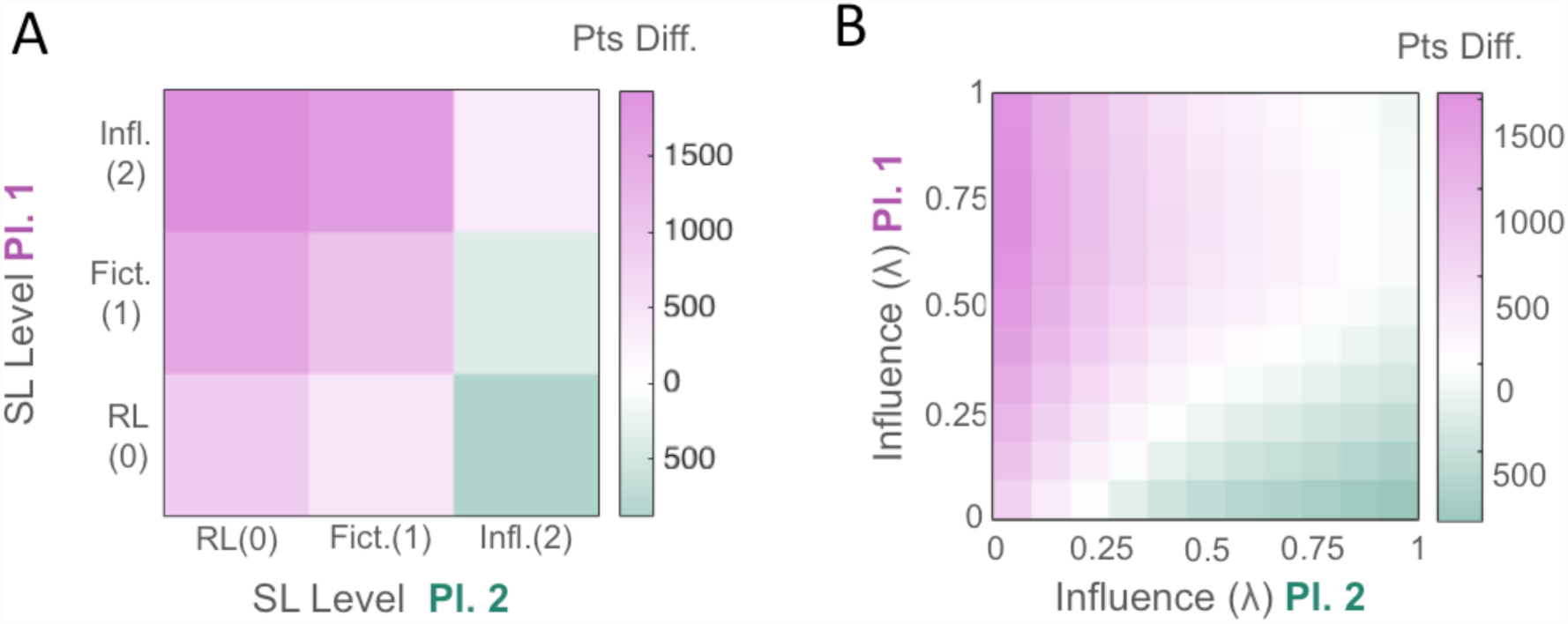
Strategic characteristic of the game: simulation of play between 2 agents varying in their Strategic Learning level (SL) shows strong asymmetry in total earnings between the 2 roles. (A) Each agent modeled by either one of the 3 models of increasing strategic complexity, or SL level (i.e. Level 0: Q-Learning, 1: Fictitious play and 2: the Influence model) played the game in one of the 2 role. Every Player1-Player2 model combination was simulated 100 times playing against each other the 100 repetitions of the game. Agents endorsing the role of Player 1 won more points on average than Players 2. In fact Player 2 agents won more that their opponent only in the situation where they were playing SL Level 2 and Player 2 agent a level below or more. (B) Agents were modelled by the Influence model varying in the values of their parameter λ from 0 (low SL level - fictitious) to 1 (high SL level - full influence), as well as their η ([0:1]). The heatmap were obtained by averaging across the whole range of η values, for each λ combination.

Altogether this preliminary analysis confirms the strategic asymmetry of our payoff matrix, revealing the strong advantage of player 1 over player 2 in the sub-optimal domain. This setting allows us to clearly test how the individual level of strategic sophistication is affected by the strategic asymmetry of the repeated competitive interaction.

### 2 Experiment 2

#### a Behavioral results

We first tested our hypothesis that our game settings triggers differences in choice behavior between the 2 roles. As predicted by our simulation analysis subjects who endorsed the role of Players 1 won more points on average than Players 2 (Block 1, B1: F(2,31)= 3.272 p= 0.0014, t(48.3)= 4.396 p<0.0001; B2: F(2,31)= 2.236 p= 0.0282, t(54.10)= 3.894 p=0.0003). In fact, across the 2 blocks, only 15% of Players 2 won more points than their opponent. The choice behavior of the 2 groups deviated on average from the optimal solution in both blocks (Player 1, P1 - B1: P(a) = 0.399(±0.065), B2: P(a) = 0.391(±0.070); P2 - B1: P(A) = 0.482(±0.098), B2: P(A) = 0.448(±0.097)) but Players 2 were the ones who deviated the most from game optimality by choosing the action “A” much more frequently than the mixed strategy equilibrium (MSNE) prescription in comparison to Players 1 (B1 t(54.09)= 3.935 p=0.0002 unequal variance, B2: t(62)= 2.696 p=0.0090). We thus aimed to test if this difference could explain the difference in performance between the 2 players. As shown on **Fig. 3**, Players 2’s deviation from MSNE was not correlated to their overall performance like Players 1 (**Fig. 3.A**), but rather to the size of their loss in the interaction (**Fig. 3.B**). In fact, the disadvantage in the interaction that was experimentally induced through the structure of the game lead Players 2 to be constrained to the loss domain, so that the closer their choice proportion was to the MSNE, the less difference in points they had with their opponent. This asymmetry in the interaction seems to have been fully exploited by Players 1 since deviation of Players 2 from the MSNE lead them to perform better than their counterpart did in this situation (**Fig. 3.C - Fig. S1.C**).

**Figure 3.**
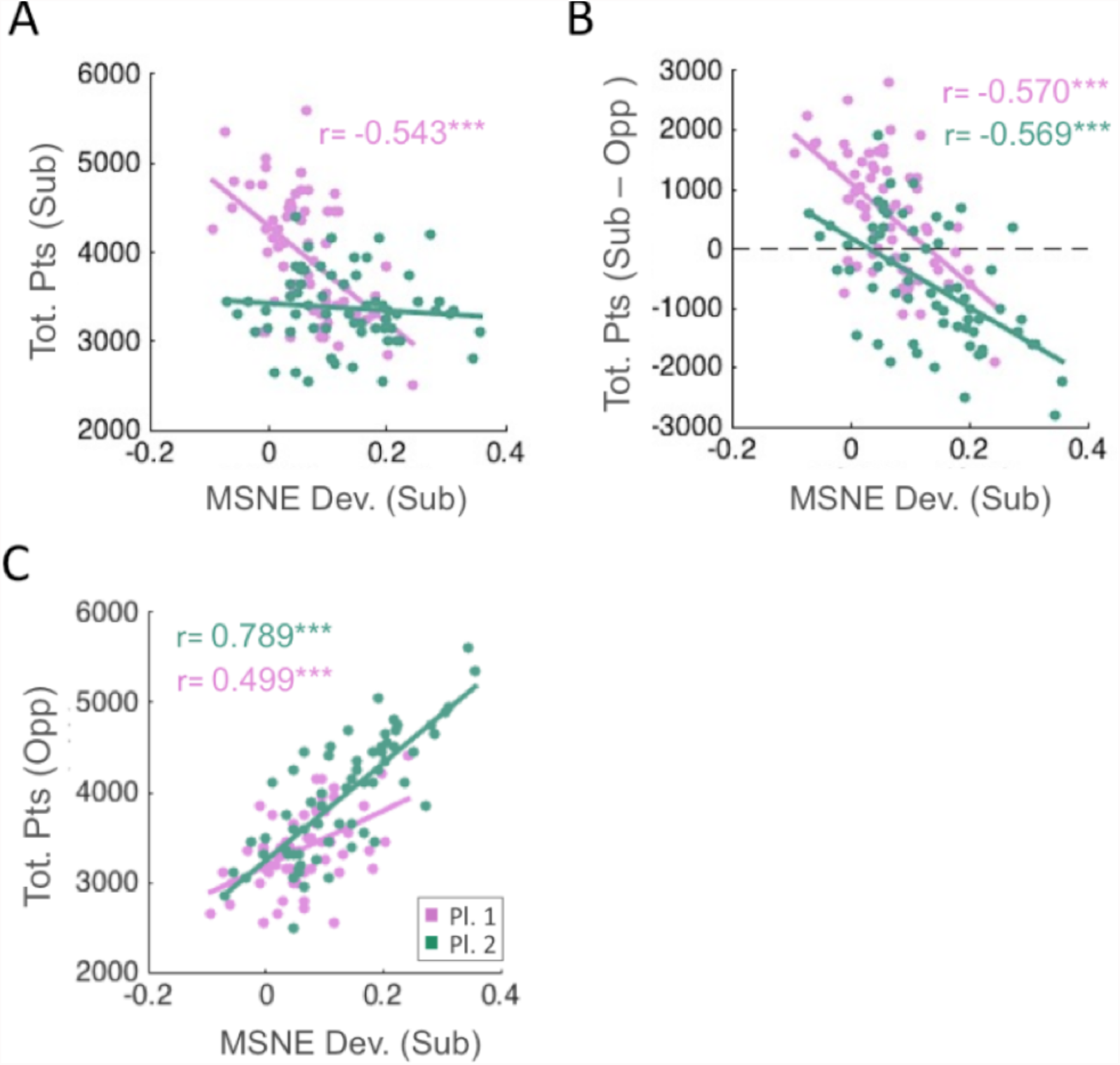
Model free analysis - Deviation from game optimality affects differently the 2 roles in the strategically asymmetric game, leading to a structural disadvantage of Players 2. (A) The closer to the MSNE the choice distribution of Players 1 is the higher their absolute performance. On the other hand Players 2 choice optimality did not lead to higher absolute performance but to a higher relative performance (B), reducing the gap in points that separate them from their opponent. This structural asymmetry lead Players 1 to fully exploit the disadvantage, their absolute performance increased more with their opponent suboptimality than Players 2 confronted to a suboptimal opponent (C).

Before investigating how the level of strategic learning affects the choice behavior of the two roles, we first tested that our prior assumption that subjects differ in their level of strategic learning was met. Our computational analysis revealed that half of our subjects behavior was best captured by the Influence model **(Fig. 4.A)**, while near one third of our population was best fitted by models of lower level of strategic complexity (less than 10% by the reinforcement learning model). Moreover, not only the subjects best fitted by the Fictitious model were also better captured by the Influence model in comparison to the reinforcement model (relative fit of the Influence) **(Fig. 4.B)**, but the better a subject’s choice behavior was captured by the high SL model, the higher the value of her Influence best fitting parameter λ was (B1: r = 0.7534, p=6.76e-13; B2: r = 0.7535, p=6.73e-13). Taken together these results reveal that the majority of subjects were engaged in some form of strategic learning throughout a gradient of strategic complexity (SL).

**Figure 4.**
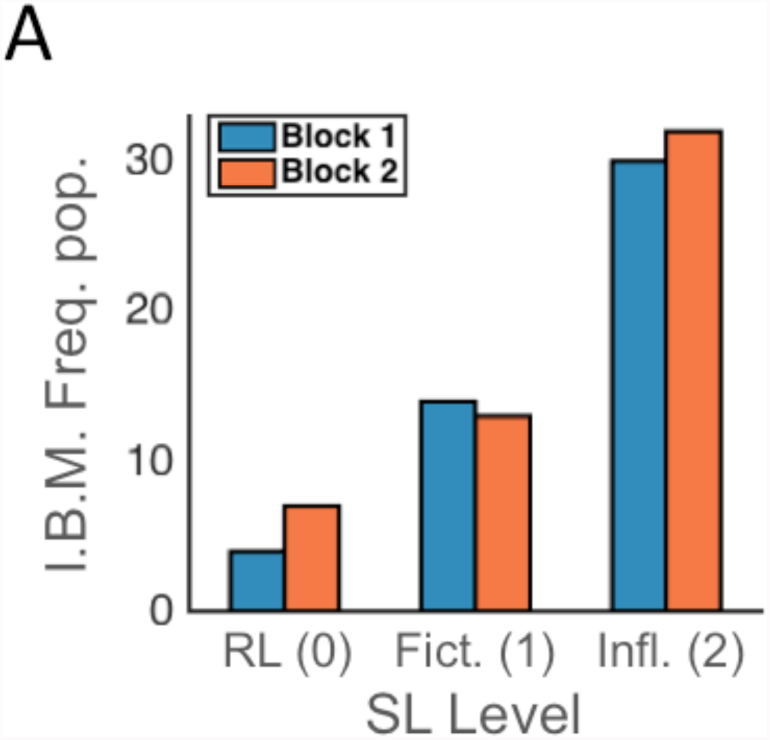
Strategic learning heterogeneity captured by our computational approach. Most of the participants engaged in Strategic Learning (SL>0). (A) Individual Best Model (I.B.M.) frequency plot. While at the population level, the Influence model fits the best the population behavior (not shown), at the individual level about half of the subjects were best fitted by high SL and one third by models of lower levels of strategic learning.

To maximize the accuracy of our individual characterization and avoid the overestimation of an individual’s strategic learning level, we conducted an extended computational analysis including additional models (see Text S1). None of the variations of the Reinforcement and Belief-Based models tested improved significantly their fit, thus confirming that most of our subjects indeed engaged in some form of strategic learning (Text S1). In fact our analysis suggests that the SL level might have been under-estimated since more than one third of the subjects previously best fitted by the Influence were better fitted by a 2nd order version of the model, which has an even higher SL level [24] **(Fig. S2)**. This nevertheless does not affect the superiority of their SL level compared to subjects best fitted by RL or Fictitious models. Although it is important to note that if the relative fit between the 2-Inf models and the simple reinforcement learning model improves the precision of the characterization of the individual SL levels, all the results presented in the following consistently hold when using as SL measure the fit of the Influence relative to the fictitious, or the value of the Influence best fitting parameter λ. Our computational analysis thus suggests an overall departure from simple reinforcement in repeated competitive interactions, with a population spread across a gradient of strategic sophistication going from value-based (Reinforcement), to low (Belief-Based) and high level (Strategic) level of learning engagement.

Besides, our simulation analysis suggested that the endogenous disadvantage of Players 2 in the game can be overcome by engaging in a higher level of sophistication than the opponent. However, the important difference in performance and game optimality observed between the two roles in our experiment lead us to hypothesize that, on average, Players 2 did not engage in a higher level of strategic learning than Players 1. Indeed, we could not reject the null-hypothesis that the two populations of SL came from the same distribution, using as SL measure either their departure from reinforcement towards models of highest strategic complexity (D(126)= 0.2031, p=0.1250; U(126)= 1682, Z = 1.7418 p= 0.0815) or the weight attributed to 2nd order belief (λ) (D(126)= 0.1719, p=0.5809; U(126)= 2036, Z = 0.0548, p= 0.9563). Note that these results hold when the 2 blocks were analyzed separately (not shown). The 2 roles did not differ either in how frequently they switched actions from one trial to the next (U(126)= 1986.5, Z =0.2913, p= 0.7708). Our results suggest that the observed disadvantage of players 2 was not due to a difference on average strategic learning sophistication but could rather be caused by a different implication of the SL level in the 2 roles.

We then investigated how the SL level of the two Players drove the dynamics of their interaction. We focus first on Players 2 behavior. The level of strategic learning engagement of Players 2 was negatively correlated with deviation from the mixed strategy equilibrium (r= −0.6455, p= 8.48e-09, SL as relative fit of the 2-Inf). Therefore, as suggested by our model-free analysis (**Fig. 5.A,B)**, their SL level was not correlated directly to the total points won in each block but to the difference in points with their opponent, so that the higher their SL level, the lower their average relative loss is (**Fig. 5.A).** Moreover, the higher their SL level was compared to their Player 1 opponent, the closer their action distribution was to the MSNE **(Fig. 5.B).** However, this was not enough to overcome the structural disadvantage and increase their absolute performance **(Fig. 5.C)**. If Players 2’s behavior seems to be constrained by their own SL level, Players 1’s behavior presents a quite opposite pattern. Their deviation from MSNE frequency was not directly driven by their SL level (r=-0.0396, p=0.7561) but by the one of their opponent (r=0.49403, p=3.34e-05), so that the higher the level of Players 2, the worse their performance was (absolute: r= - 0.5826, p= 4.40e-07 and relative: r= −0.4650, p= 0.0001). Since Players 2 who engaged in a higher SL level deviated more from the High Reward action to play closer to the MSNE, they pushed Players 1 to adapt by engaging in a higher SL level. Indeed, given the structural advantage they had in the game, the better they were at anticipating their opponent’s behavior (higher SL level than their opponent), the higher were their relative earnings (r= 0.4380, p= 0.0003) and their absolute performance (**Fig. 5.C**). These results hold when comparing these behavioral measures between high and low SL level (median split) in the 2 roles.

**Figure 5.**
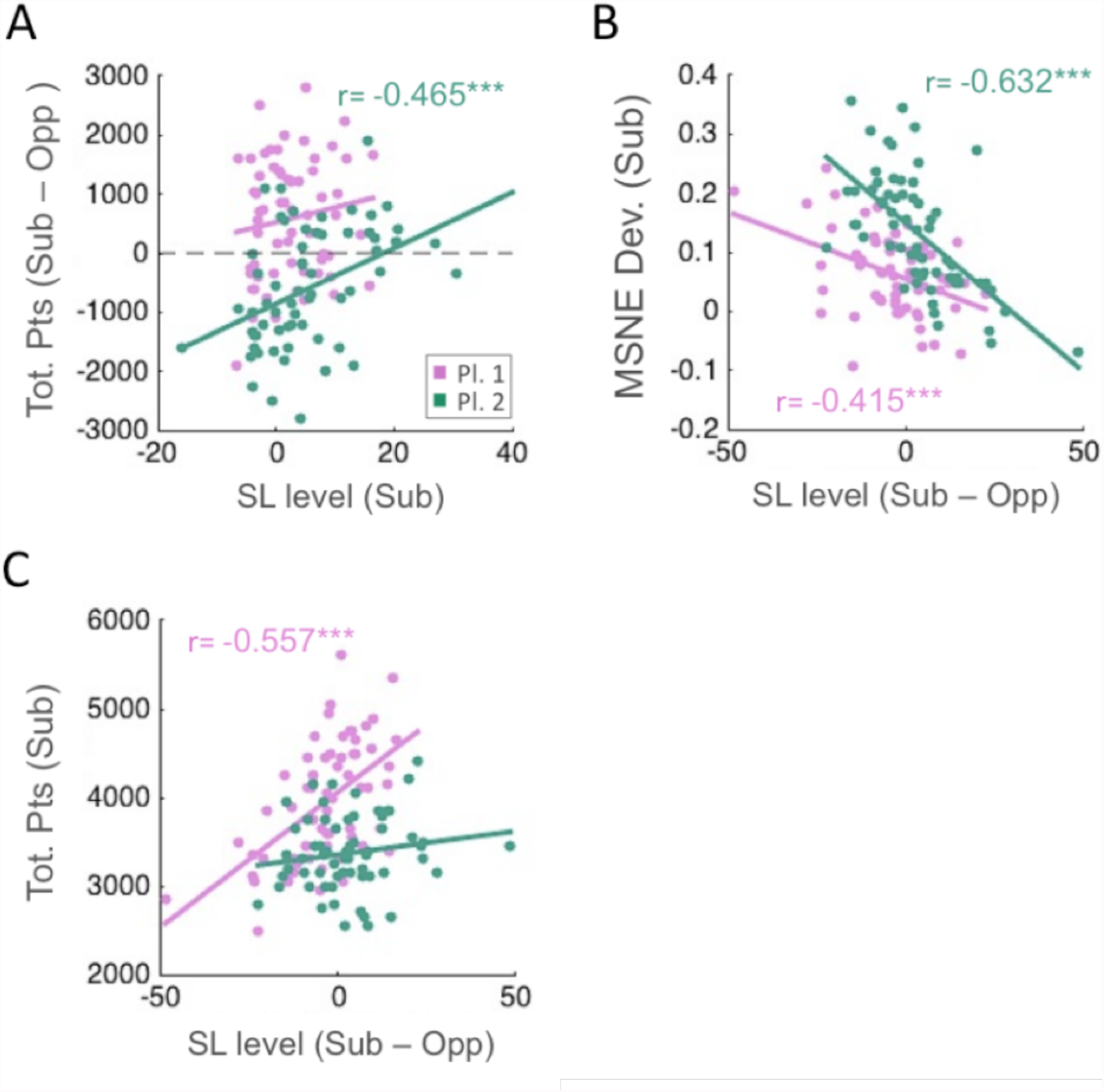
Model-based analysis - The Strategic Learning level (SL) of the Players 2 in the game, conditions their capacity to overcome the structural disadvantage of their position in the game. (A) The higher the SL level, measured here by the difference in the fit between the (second order) Influence model compared to the fit of the RL^†^, the more Players 2 reduced their disadvantage compared to their opponent. (B) The higher their SL level compared to their opponent the closer to the MSNE they played. In fact both role converge towards the equilibrium distribution, only Players 2 tend to deviate much more when not engaging in strategic learning. (C) Albeit decreasing the gap in points with their opponent Players 2 could not on average increase their overall performance, constrained by both the structure of the interaction and their own SL level. (^†^ Similar results were obtained when running the correlation test analysis with the relative fit of the first order Influence. Using the Influence parameter λ values as measure of the SL level or comparing Low and High SL IBM groups of subjects lead conserved the main statistical effects.)

To capture the simultaneous effects of both the subjects’ and their opponent’s SL level on the subject’s own choice behavior, we ran 3 GLM analyses that confirmed that Players 2’s behavior was impacted mainly by their own level of strategic learning sophistication, and Players 1’s behavior mainly by the SL level of their opponent (**Fig. S3.A.B).**

This dynamic can be further unfolded by looking at the choice accuracy of the subjects. Players 2 who engaged in higher SL level managed to overrule the value-based sub-optimal bias towards the high reward action. Instead of repeatedly selecting action A, easily predictable by their opponent in the advantageous situation, they switched action more often from one trial to the other (r= 0.3419, p=0.0057), so that they got more frequently the high reward when they chose action A **(Fig. S4.C)**. Conversely this lead Players 1 to compensate, to avoid deviating more from the optimal play, by engaging in higher strategic learning eventually leading to also increase their accuracy **(Fig. S4.C)**.

This overriding of the prime tendency for Players 2 to go for the high reward by engaging in higher level of strategic learning level was also observed from one choice to another. Using a logistic regression analysis we can take a closer look to the series of choices to investigate how the previous actions impact the current decision (details provided in **Text S1**). This analysis revealed that on average subjects consistently alternated their choices every 2 trials independently of their role (**Fig. S5.A)**, but that only Players 2 tended to persist in selecting the action linked to the high reward, taking less into account the opponent’s last choice (**Fig. S5.A,B)**. And the more Players 2 engaged in strategic learning the more they would alternate their choice (**Fig. S5.C)**.

Altogether, our analyses suggest that among the subjects endorsing the role of Player 2 in this experiment, only the ones who had a high level of strategic learning sophistication could detach from the game sub-optimal focal point to overcome the structural disadvantage they had in the game interaction. Their opponent, albeit in the easy position, was then forced to adapt and at the end to follow the Players 2’s choices to avoid as much as possible to lose their advantage. The leader becomes the follower. This hypothesis was further backed up by our initial simulation. Indeed, running the same type of analysis on our simulation results leads to a very similar dissymmetry in the implication of the SL level between the advantageous and the disadvantageous role **(Fig. S6).**

#### b Correlation with additional cognitive tasks

Our main hypothesis was that Players 2’s disadvantage in the game would push them to make the effort to use their strategic thinking abilities, so that their performance in the game, as measured in terms of heterogeneity of the SL level of the computational model that best accounted for each subject’s behavior, would be more stable across opponents and rely more on cognitive constraints than it would be the case for Players 1. Conversely we hypothesized that Players 1’s strategic learning engagement would be influenced more by individual traits and problem-solving or planning abilities, because their advantage in the game would leave them free to play with the SL level they are habituated to use outside the laboratory. To test our hypothesis we first looked at the consistency of the SL level of Players 2 across blocks compared to Players 1. We found that their SL level was significantly more consistent (89% of the subjects were best fitted by the same class - low / high - of SL level models in Block 1 and Block 2) compared to Players 1 (59%, Fisher exact test: N = 49, p = 0.0269). Also the correlation in SL level across the 2 blocks was significant for Players 2 only (P1: r = −0.0781, p= 0.6710; P2: r = 0.7171, p=3.88e-06). Moreover, no difference on average (or distribution of) SL nor choice behavior (deviation from MSNE, absolute or relative performance, choice accuracy of high reward action, frequency of switch) was found between the 2 blocks for each role. Interestingly, our analysis nevertheless revealed that Players 1 chose faster in the second trial block (P1: t(62)= 2.7102, p = 0.0087, P2: t(62)= 1.9009, p = 0.0620), suggesting some adaptation of Players 1’s play in the second block.

Finally, we compared the SL level of the subjects and their individual performances in the additional tasks and questionnaires. At the population level only the CRT score (used as a proxy of reasoning ability in the literature) was higher for high SL vs. low SL (median split: U(62)=313.5,Z= 2.7678, p=0.0056; r=0.2572, p=0.0402) and in Block 1 only. When comparing the performance in the additional tasks in high vs. low SL level (median split) subjects for each role separately, we found that high SL Players 1 in Block 1 only had a higher CRT score (U(30)=54, Z=2.8891, p=0.0038), performed better in the Raven test (U(30)=62, Z=2.5389, p=0.0111), and were on average more successful in the Tower of London task (ToL: t(25)=2.0675, p=0.0491; ToL (difficult condition: high Goal Hierarchy, high Search Depth): U(25)=46, Z=2.3271, p=0.0199). No correlation between the performance in any the additional tasks was found with the SL level of Players 2 in any of the 2 blocks. No difference was found between subjects based on the role they endorsed in the game in terms of demographics (Age, salary and education level) nor additional cognitive tasks performances (Working Memory, CRT, Raven, ToL). No difference either in performance in these additional tasks was found between subjects who were consistently fitted by the same SL model in both blocks and the one who switched SL level between the two.

Our results suggest that regardless of the role, the subject’s level of strategic sophistication does not depend on the level of the opponent nor the role endorsed in the game but might be related more to different individual cognitive abilities: subjects playing as Player 2 seem to be limited in their propensity to engage in strategic learning preventing them to fully compensate their disadvantage in the game interaction, while the heterogeneity in SL level observed in Players 1, already in a dominant position, might be more driven initially by their executive cognitive abilities (problem-solving, planning). If this is true then Players 2 ability to engage in strategic learning might be correlated to their ability to reason strategically (i.e. in an iterative fashion).

During our experimental session, subjects were provided with a second task meant to test specifically the hypothesis that the level of strategic learning engagement in the repeated game matches the ability to play the Nash equilibrium in one-shot games [26]. This task was composed of different types of one-shot games played 8 times each, in a random order, with no feedback. In one type of games (Dominant Solvable Other, DSO), higher strategic sophistication was required to form correct belief over the opponent’s action and best respond to it but not in the other (Dominant Solvable Self, DSS). The analysis detailed in the **Text S1** did not allow us to reject our null-hypothesis of an absence of direct mapping between strategic learning and strategic reasoning at the population level. This therefore suggests that different cognitive processes are involved in the engagement in strategic sophisticated play in a repeated game interaction with feedback and in static one-shot games without feedback. However we found that the more Players 2 reached the N.E. in DSO (requiring higher level of strategic sophistication) the closer to the Nash their frequency of action(a) was in the first Block in the repeated game (all trials B1: r=-0.3822 p=0.0309; B2: r=-0.3078 p=0.0865). This correlation, specific to Players 2, was the strongest in the first trials of the all repeated game experiment (B1 t(1:10): r=-0.5647 p=7.6e-4 - not sig. for following bins; B2 t(1:10): r= −0.1992 p= 0.2743 - not sig. for following bins). Using 2 other, more precise, measures of strategic reasoning developed in SI, lead to similar results (SR: r=-0.5034, p= 0.0063; SR’: r= −0.6853, p=0.0068), no correlation was found with the % of NE in DSS.

Taken together these results suggest a transition from static strategic reasoning to on-line computation and update of beliefs over the opponent’s behavior when sufficient choice outcomes are observed.

### 3 Experiment 3

To better characterize the interplay of strategic learning sophistication with the strategic asymmetry in game interaction, we conducted a second study with human subjects in which we controlled for the opponent’s behavior by making the subject play against a computer opponent (instructed) and specifically manipulating the SL level of this opponent. The goal of this experiment was to replicate our initial results and test the specific hypotheses derived from them, that subjects endorsing the disadvantageous role in this strategically asymmetric game were constrained in their choice behavior by their own SL level, while subjects playing in the advantageous position will indeed have the strategic space to adapt, given their SL level, to the behavior of their opponent.

Our Human vs. Agent design allowed us to test the 2 main hypotheses from Exp. 1 and 2: 1) that the role impacts the average performance and game optimality (Players 1 perform better than Players 2), but not the overall SL distribution of the two groups. 2) that Players 1’s choice behavior should be impacted by the identity of the opponent, with lower performance against High SL agents (Influence) compared to Low SL agents (fictitious). Players 2’s behavior on the other hand should only be affected by their own SL level, not the opponent. Beforehand we simulated once again the experiment by making agents with different SL level play against the 2 algorithms. Our results show the selective effect of the opponent SL level manipulation on the Players 1 we expect to see in the actual experiment, thus confirming the adequacy of our design (**Fig. S7).**

On average, Players 1 won more points (t(142)= 8.5298 p=2.79e-14) and had a distribution of choices closer to the Mixed Nash Equilibrium (t(142)= −4.5144 p=1.32e-05 unequal variance) than Players 2. Our Model-based analysis replicated nicely the distribution and SL gradient across the population observed in Exp.1 (**Fig. S8)**. And, as in Exp.1, no difference in SL distribution was found between the 2 roles. But this time participants were playing against an algorithm, not against another human. And since they played the repeated game in the same experimental conditions as in Experiment 1, we tested if this difference affected their behavior. For each of the 2 roles, no significant difference was found in performance (total points, points difference with the opponent) between the two experiments. However, a trend towards higher strategic learning engagement when playing against algorithms was observed. When comparing low vs. high SL (median split), Players 1 engaged in strategic learning were found to have a higher SL level in the second experiment (rel. fit 2-Inf: U(66)=233, Z=4.2082, p= 2.57e-05, λ parameter: U(66)=368, Z=2.5495, p= 0.0108 - similar results were obtained when comparing the SL level between subjects best fitted by the Influence models).

No difference in mean SL level (nor distribution) was found between the 2 roles. Running an ANOVA to test if the SL level was modulated by the opponent encountered did not result in any significant effect either. As in experiment 1, Players 2 were most consistently fitted by same SL level models between the 2 opponents than Players 1 (P2= 0.84 prop. same low/high SL: P2=0.84, P1 = 0.57; Fisher exact test: N = 58, p = 0.0259).

We next tested our second hypothesis regarding the specific effect of the opponent on the choice behavior of the subject given the role endorsed in the experiment. As shown in **Fig.6**, only Players 1 were affected by the identity of the opponent, exactly as predicted by the simulation (**Fig.5.A.B, Fig.S7.A.B)**.

**Figure 6.**
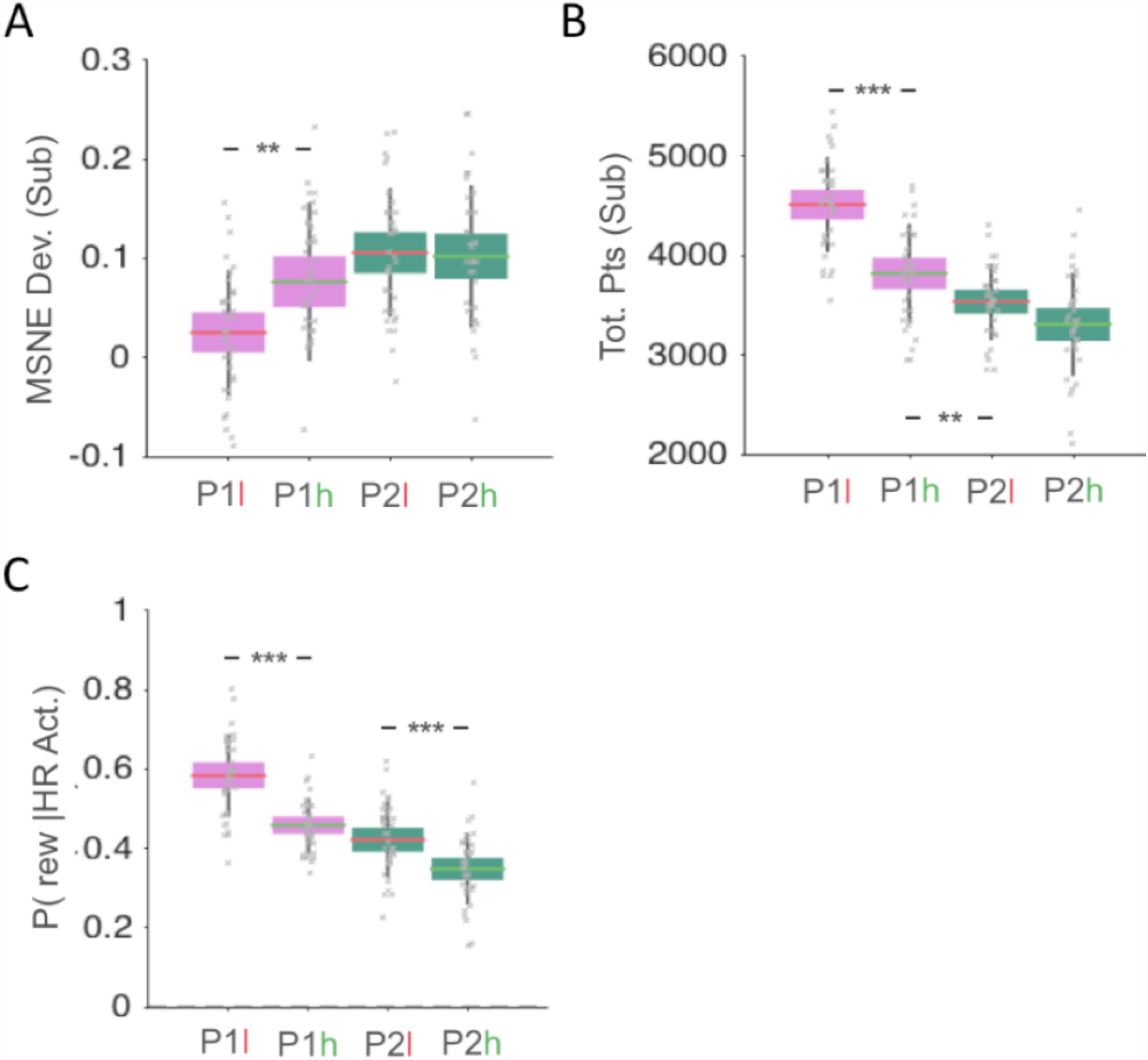
As hypothesized, only the subjects endorsing the role of Player 1 (pink) in the repeated game were affected in their choice behavior by the SL level of the (computerized) opponent encountered. (A) Players 1 frequency of choice is closer to the MSNE distribution when playing against the low SL opponent compared to the high level. No difference in percentage of deviation from MSNE distribution (p(a)=1/3) was found between the 2 opponent’s block for Players 2. (B) When opposed to the low SL level opponent, Players 1 won on average more points in total than when playing against the high SL. No difference was found for Players 2. (C) When opposed to the low SL opponent both Players, 1 and 2, were more frequently rewarded when playing the high payoff action (b for player 1 and A for player 2), a proxy for choice accuracy, compared to the high SL level. Effect size was however higher for Player 1 (Cohen’s d= 1.1909) than Player 2 (d= 0.7633).

To refine our analysis, we ran the 3 GLMs we used in Exp. 1, taking into account not only the level of the computerized opponent (low or high) but also the SL level of the subjects. Our results replicate strongly the asymmetry found in our first experiment and observed in the average results (**Fig.6)**: Subjects playing as Player 1 have their deviation from MSNE as well as their performance affected by the level of the opponent; Conversely Players 2’s choice behavior is modulated only by their own Strategic Learning level (**Fig.6)**. In this experiment however, Players 1’s behavior seems to have been influenced not only by the level of the opponent but also by their own strategic learning engagement (**Fig.S9)**. This effect could be due to the constrain our design added on their opponent’s behavior.

At the end of the experiment subjects were provided with an additional task aimed at capturing more precisely the working-memory capacity of our population (namely 2 and 3-Back tasks - see Text S1 for details). We observed a trend towards a higher performance and RT in this task for high SL Players 2 (median split) only when playing against the high SL opponent (% correct in 2-Back: t(33) = 2.2047 p=0.0361, 3-Back: t(33)=1.7420 p=0.0908, albeit a higher % for High SL =75.4(±9.5) vs. low SL=69.9(±8.9), reaction time 3-Back: t(33)=2.2999 p=0.0279). Albeit weak, this effect suggests that the subjects playing in the disadvantageous role might be cognitively constrained in their higher engagement in strategic learning when confronted to a highly sophisticated learner.

## VI DISCUSSION

The present study aimed at testing the prediction that the structure of a repeated game interaction can lead to strategic asymmetry depending on the way it facilitates the engagement in sophisticated learning. More precisely, the hypothesis was that a dissymmetry in the overlap between reward structure and MSNE (even when there is still symmetry in maximum possible payoffs between players) can differently engage human subjects in using sophisticated strategic learning so that their overall performance does not always reflect an individual’s general ability of strategic learning or strategic reasoning.

This hypothesis was rooted in research in behavioral economics and cognitive sciences suggesting that humans can use information available about their counterpart to form beliefs over their intentions [3]. Indeed, in the case of repeated games, where social interactions are reduced to actions and rewarded feedbacks, payoffs convey informational value about the opponent’s behavior, and beliefs become analogous to a mapping of action-outcome contingencies, as suggested by the model-based reinforcement learning framework [20]. Recent studies suggest that similar brain computations might be involved in the decrease of the uncertainty (increased prediction) over the opponent’s next choice allowing one to maximize her overall outcome of the interaction (best response to beliefs) [27]. Moreover, inferential processes might be implicated in the iterative incorporation of the strategic nature of the interaction, not only considering one’s behavior but the interplay of past actions in the history of play, ultimately increasing belief accuracy over an opponent also capable of belief-learning [26], [32]. Nevertheless, important heterogeneity has been observed in the level of engagement in such high-order belief (strategic) learning among individuals [24], [25].

We thus hypothesized that the reward structure of the interaction might affect subjects differently given their capacity to engage in strategic learning, depending on how much best-response to reward-based and belief-based learning overlap. Based on this prediction we developed a 2×2 strategically asymmetric game where the two roles were meant equal (same payoff distribution and expected payoff at MSNE), but in which inequity arises among individuals with different SL levels, from the discrepancy in one role (disadvantaged position) only between the highest reward action (focal point) and the MSNE distribution.

To test this hypothesis, we combined agent simulation and behavioral experiments, the latters being either unconstrained (human-human interaction) or constrained (human-computer). Our two behavioral experiments lead to the same conclusions, predicted by our simulation analysis. First, at the population level, subjects endorsing the disadvantageous position during the repeated game interaction earn significantly less than their opponent. Their choice distribution also deviated more from the MSNE prescription. Second, the more the disadvantaged participants engaged in strategic learning, the more they overcame the strategic asymmetry. This effect was even stronger when the opponent did not fully engage in belief-learning. Forming accurate beliefs over their opponent allows these subjects to reduce their disadvantage in total earnings and to play closer to the MSNE. Conversely, the choice optimality and therefore the absolute performance of the participants playing in the advantaged role was modulated only by the behavior, and ultimately the SL level, of their opponent, but not by their own capacity to engage in strategic learning.

These results provide clear evidence for sophisticated learning in repeated interactions [4], [33] and the central role of belief accuracy in equilibrium play [34], [15]. Moreover, our study shows how the reward structure of the repeated game interacts with the observed heterogeneity in belief-learning at the population level by creating a tension between rewards and beliefs. Empirically, we show that the high reward attracts maximizing behavior and creates a focal point, which can easily be exploited by a low strategic learning opponent [35] when not aligned with the MSNE prescription. In our stage game the two players had identical expected payoffs. But the fact that for only one of the two roles the focal point corresponded to the action that was theoretically the most optimal to select, creates an endogenous asymmetry, strategic in nature, which reveals itself throughout the interaction.

While Mixed Strategy Nash Equilibrium (MSNE) theoretically prescribes a probability distribution over possible actions that ensures to each involved agent that they would have no incentive to deviate if they all follow it, in practice, however, humans have been found to typically deviate from this equilibrium [6]. In repeated games, patterns of aggregated choices have been sometimes found to converge towards MSNE [8] while other studies have found a strong departure from MSNE [7]. Importantly, following MSNE requires for each subject to show some level of randomness in their behavior. This enables subjects in practice to approximately stick to the action probability distribution prescribed by the MSNE without displaying a trivial repetitive behavior which would have entailed the risk to be detected/predictable by the opponent. This empirical finding is surprising as humans have been systematically proved to be bad randomizers [9]. Indeed, serial dependency in between actions is usually observed [10], leading authors to suggest that learning might lead to MSNE [11], [12]. Here the employed asymmetric structure of our game enabled to bring further insight about some of the conditions in which humans can deviate from MSNE. Introducing a focal point in the game payoff matrix pushed disadvantaged players to deviate from MSNE, while forming accurate beliefs over their opponent allowed these subjects to reduce their disadvantage in total earnings and to play closer to the MSNE.

Previous studies also used a computational approach to capture as precisely as possible the choice behavior of humans in repeated 2×2 games [28], [36], [37]. We went a step further and manipulated the interaction structure to show that the level of the strategic sophistication of individual’s learning drives the formation of higher order beliefs and allows them to disengage from the attractions of immediate outcomes and move closer to optimality.

Crucially, no correlation was found in any of the 2 experiments between the SL level of the subjects, in any of the two roles, and the one of the opponent. This result replicates the correlation in strategic learning level found in previous studies in which humans were confronted to different opponents also varying in their SL level [24], [38]. It is worth noting, however, that recent research suggests that arbitration between model-free and model-based learning can be affected by the volatility of the environment [39]. Indeed, in our two experiments most of the subjects did engage in a rudimentary form of belief-learning which does not reject the hypothesis that parts of a subject’s learning mechanisms may be with low sophistication (model-free). Moreover, our computational approach was meant to measure the overall level of strategic learning sophistication embedded in individual choice series, and does not allow to track local changes in strategy. In fact, as previously observed [40], [41], [42], the SL level of the opponent strongly impacted the behavior of the subjects interacting in the advantageous position. However, in our study the influence of these subjects’ own SL level on their choice behavior was reported as weaker than the influence of their opponent. One hypothesis for this result is that the sophistication of their beliefs did not condition their behavior, which obviously comes in contradiction with the above-cited literature. An alternative hypothesis is that the accuracy of their beliefs was already sufficient to maximize (up to a certain individual threshold) their earnings, and that engaging in higher level of strategic learning did not present a net advantage. We argue that the present study provides the first experimental evidence in favor of the latter explanation. First subjects playing in the advantageous role won on average more points throughout the interaction than their opponent, even when opposed to a high SL computerized agent (t(70)=2.7833, p=0.0069, **Fig.6.B).** Second, albeit good performance, these subjects presented low consistency in SL level across interactions in both experiments. Third, their behavior was correlated to planning and problem-solving scores captured in additional tasks in the first interaction block only, while in the second block conjunctional evidence of behavioral re-adjustment (faster choices for no change in performance) were observed.

In contrast, disadvantaged subjects behavior were found to be solely conditioned by their own level of strategic engagement, not the one of their opponent. They also presented much higher consistency across interactions, and evidence of a correlation between working-memory and their SL level was found. Crucially, the role endorsed in the repeated game did not seem to impact the SL level of the subjects in either of our experiments.

Altogether, our results suggest a dissociation between a strategic learning engagement bounded by individual cognition for subjects endorsing the disadvantaged position, and what has been proposed to resemble a cost-benefit analysis process [43] for the subjects ensured to dominate the interaction at lower SL level. This distinction between bounded cognition and bounded rationality in suboptimal play has also been observed in static games by Friedenberg et al [31].

In behavioral game theory the concept of bounded rationality broadly assumes that the capacity of the agents to grasp and use all the required information leading to equilibrium are somehow constrained [44]. In this line, a theoretical framework which has accumulated growing support in the past decade has been proposed to explain deviations from optimal choices in static games: level-k models [15]. This class of model relaxes the assumption of full rationality and assumes that players actually best respond to incomplete beliefs varying among individuals in their degree of sophistication (k) over the behavior of their opponent, themselves considered as only capable of a lower order of beliefs (k-1 or <1, see [45]). This hierarchical organization of beliefs is very close to the computational framework previously developed in [25], and that we used in this study. In our approach, strategic learning sophistication (SL) is modeled as a hierarchy of different levels of computations: SL0 corresponds to reinforcement learning which computes action values based on the past reward experienced and is agnostic about the choice behavior of the opponent; SL1 is modeled as a fictitious play which best responds to the opponent’s probability distribution computed from its past choices; SL2 is modeled as an influence learning process which assumes that the opponent is also learning in a way analog to a fictitious play and thus that its own past actions can have an influence over the action probability of the other (we also included a SL2+ learning rule that considers that the opponent is also learning through influence). Based on this correspondence between the 2 classes of models, we have hypothesized that a direct mapping might exist between the level k in static games and the SL level in repeated interactions [26]. We tested this hypothesis but failed to reject a direct correlation between the two measures of strategic sophistication (see **Text S1**). Nevertheless, we observed that for the subjects endorsing the disadvantaged position in the repeated game, a correlation was found between their strategic reasoning (level k) and the frequency of play close to the MSNE in the very first trials of the first interaction. This result therefore suggests another type of relationship between the strategic reasoning and strategic learning models of bounded rationality: at first, when beliefs cannot be anchored in enough observations, subjects with a strong incentive to take over the interaction are guided by their ability to reason in an iterative fashion (level k); but when enough experience is accumulated, subjects with the capacity to engage in strategic learning form and update beliefs as accurate as possible over their opponent’s behavior. This hypothesis appears promising to us since it echoes other research on the influence of priors in social inference [46], and therefore calls for proper testing in laboratory.

It is worth noting that another source of sub-optimality has been suggested in the behavioral economics literature: heterogeneity in best response. It has been proposed that social preferences for instance could bend utility functions [47]. If our study did not allow to directly test this hypothesis, we still observed a higher strategic engagement in advantaged subjects capable of high strategic learning when confronted to an algorithm compared to another human (social framing effect as in [24]). This result might suggest that social preferences such as altruism or sensitivity to inequity could be reflected in a lower exploitation of their advantageous position when playing with a human counterpart [48].

Altogether, our results reveal three possible sources of variability in strategic behavior during repeated game interactions, which makes this behavior not simply constrained by the subjects’ cognitive capacities only, and can thus help reconcile previous contradictory findings that subjects’ strategic behavioral performance can be either predicted by their cognitive capacity [49], [50] or not [24]. The first source of variability, that we call exogenous, is driven by external factors such as the payoff structure, the salience of the different outcomes of the game, and also the prior knowledge over the opponent. A second source of heterogeneity in game play is endogenous, with differences in social preferences but also motivation [51] or sensitivity to rewards [52]. Finally, a third type of variance emerges from the two previous ones along experience with the repeated game, leading to specific dynamics of repeated choice behavior. Indeed, even if our experimental setting did not allow further investigation of the phenomenon, it seems clear that the SL level of subjects in the disadvantageous position drove the interaction, while their opponent in the dominant position would simply track and adapt to changes in behavior and ultimately followed them. Leader-follower dynamics have been observed in repeated games [53]. However, a precise understanding of the underlying behavioral forces remain unclear [54]. Predicting the learning dynamics in play by tuning the structure of the interaction can help study critical behavior such as strategic teaching [55].

The present study brings further support to the pertinence of the cognitive neuroscience framework of learning for the analysis of repeated non-cooperative game behavior. Moreover, we advocate in favor of the use of model simulations in the field, that 1) allow to take the most of a normative framework to optimize experimental design and make precise predictions regarding the expected results [56] and 2) open the possibility to refine agent-based simulation analyses in order to better characterize the interplay between the level of strategic learning and the structure of the game underlying the repeated interaction.

At a broader scope, this study points in the direction of a systematic consideration of between-subjects differences and the interaction effect between human variance in learning and the way strategic interactions are constrained. Taking into consideration asymmetric facilitation can help study the emergence of social hierarchy and strategic dominance in interactions [57], but also better understand how inequity arising from the interaction between the environment and endogenous differences could be reduced in real-life social interactions [58].

## ACKNOWLEDGMENTS

We would like to thank Jan Drugowitsch for letting us use his matlab toolbox of optimization through slice sampling.

## SUPPORTING INFORMATION

### Text S1

This is a document containing supporting information regarding models, statistical methods, experimental details, additional data analyses.

